# Avoidance response to CO_2_ in the lateral horn

**DOI:** 10.1101/342675

**Authors:** Nélia Varela, Miguel Gaspar, Sophie Dias, Maria Luísa Vasconcelos

**Affiliations:** Champalimaud Research, Champalimaud Centre for the Unknown, 1400-038 Lisbon, Portugal

## Abstract

In flies, the olfactory information is carried from the first relay in the brain, the antennal lobe, to the mushroom body (MB) and the lateral horn (LH). Olfactory associations are formed in the MB. The LH was ascribed a role in innate responses based on the stereotyped connectivity with the antennal lobe, stereotyped physiological responses to odors and MB silencing experiments. Direct evidence for the functional role of the LH is still missing. Here we investigate the behavioral role of the LH neurons directly, using the CO_2_ response as a paradigm. Our results show the involvement of the LH in innate responses. Specifically, we demonstrate that activity in two sets of neurons is required for the full behavioral response to CO_2_. Using calcium imaging we observe that the two sets of neurons respond to CO_2_ in different manners. Using independent manipulation and recording of the two sets of neurons we find that the one that projects to the SIP also outputs to the local neurons within the LH. The design of simultaneous output at the LH and the SIP, an output of the MB, allows for coordination between innate and learned responses.

## INTRODUCTION

Animals use the olfactory system to find partners or food and to avoid predators. To a certain extent the ability to navigate the olfactory environment is hardwired. This ability is expanded with life experiences that result in olfactory associations. The architecture of the olfactory system is comprehensively characterized in the fruit fly and it is remarkably similar to the mammalian olfactory system (1,2).

Olfactory sensory neurons express a single odorant receptor. Olfactory sensory expressing the same receptor project to the same glomerulus in the first olfactory center in the brain called antennal lobe in the fly. Most projection neurons innervate a single glomerulus and carry the information to higher brain centers: the mushroom body (MB) and the lateral horn (LH) (3,4). The MB is critical for olfactory associations (5). The LH was ascribed the role of innate response based on MB silencing experiments (6,7). Projection neurons from the antennal lobe connect to the MB without apparent spatial selection whereas at the LH axonal arbors from different glomeruli interdigitate in a stereotyped fashion (3,4,8–11). The stereotypy is consistent with the proposed role for the LH as the center for innate olfactory processing. Though the axonal arbors of projection neurons interdigitate the findings that the arbors of projection neurons that carry information on pheromone and food odors segregate within the LH and that a region within the LH is tuned to repulsive odors suggest dedicated processing areas within the LH (12,13). Lateral horn neurons (LHNs) that respond to the male pheromone 11-cis-vaccenyl acetate were identified based on the expression of the male-specific form of the transcription factor fruitless (14,15). One cluster of male LHNs responds specifically to the pheromone. These results open the possibility that each odor has a cognate LHN. Indeed, a theoretical study supports this connectivity (16). However, activity and anatomy of other LHN clusters suggests a mixed model of connectivity (17). While connectivity at the LH is being scrutinized, direct evidence for the functional role of the LH is still missing.

One of the innate responses with the highest magnitude is the response to CO_2_. Unlike most insects *Drosophila melanogaster* avoids CO_2_ when tested on a T-maze. The aversive response up to 2% CO_2_ is solely mediated by antennal neurons expressing the CO_2_ receptor complex GR21a-GR63a which connect to the V glomerulus in the antennal lobe (18,19). Synaptic inhibition of GR21a-GR63a expressing neurons abolishes the avoidance response to low concentrations of CO_2_ (20). Conversely, artificial stimulation of CO_2_ sensing neurons with light elicits the avoidance behavior typically observed in response to CO_2_ (21). Not all responses to CO_2_ are mediated by GR21a-GR63a neurons. Higher CO_2_ concentrations elicit a response to acid which is processed in a separate glomerulus (22). Also, a recent study shows that the response to CO_2_ is state-dependent with high activity flies moving towards CO_2_ and low activity flies avoiding CO_2_ (23). The attractive response does not require GR21a-GR63a receptors, instead it is mediated by the ionotropic co-receptor IR25a.

Here we address directly the behavioral role of the LH neurons using the CO_2_ avoidance to low concentrations to probe the requirement of the LH for innate responses. We demonstrate that activity in two sets of neurons is required selectively in the behavioral response to CO_2_.

## RESULTS

### Neurons labeled by lines *R21G11* and *R23C09* process CO_2_ avoidance

We chose to investigate the role of LH neurons in the context of the response to CO_2_ due to the high magnitude of the innate response. To identify neurons involved in CO_2_ avoidance we performed an inhibitory screen of fly lines labeling LH neurons (Figure S1). Through visual inspection of the expression pattern of Janelia’s collection of *GAL4* lines we selected 32 lines with obvious LH innervation (24,25). To silence the neurons we expressed the inwardly rectifier potassium channel, *Kir2.1* (26), that hyperpolarizes neurons and thus decreases the probability of firing an action potential. In the screen and in other behavioral experiments with *GAL4* lines, we restricted *Kir2.1* expression to adult stage by using a temperature sensitive *GAL80 (GAL80^ts^*, see materials and methods) (27). The 32 lines were tested on a T-maze where flies were allowed to choose between air and 0.5% CO_2_ (Figure S1A). Eight lines showed a significant reduction in avoidance (multiple t-test corrected with Holm-Sidak method, p<0.05,) and when retested, three of them exhibited a consistent reduction in avoidance to CO_2_ (Figure S1B). Line *R65D12* was discarded due to innervation in the V-glomerulus (data not shown). Neurons in lines *R21G11* and *R23CO9* (which we will henceforth call *21G11* and *23CO9* neurons) are necessary for the behavioral response to CO_2_ (Figure 1A, here tested to 1% CO_2_). Since the requirement of the V-glomerulus bilateral projection neurons and the MB for CO_2_ avoidance is feeding state dependent (10), we tested if feeding state also affects the contribution of *21G11* and *23C09* neurons in the avoidance response of the fly. We observe that starvation does not alter the phenotype, indicating that the involvement of *21G11* and *23C09* neurons in CO_2_ response is independent of the feeding state of the fly (Figure S2A).

**Figure 1.**
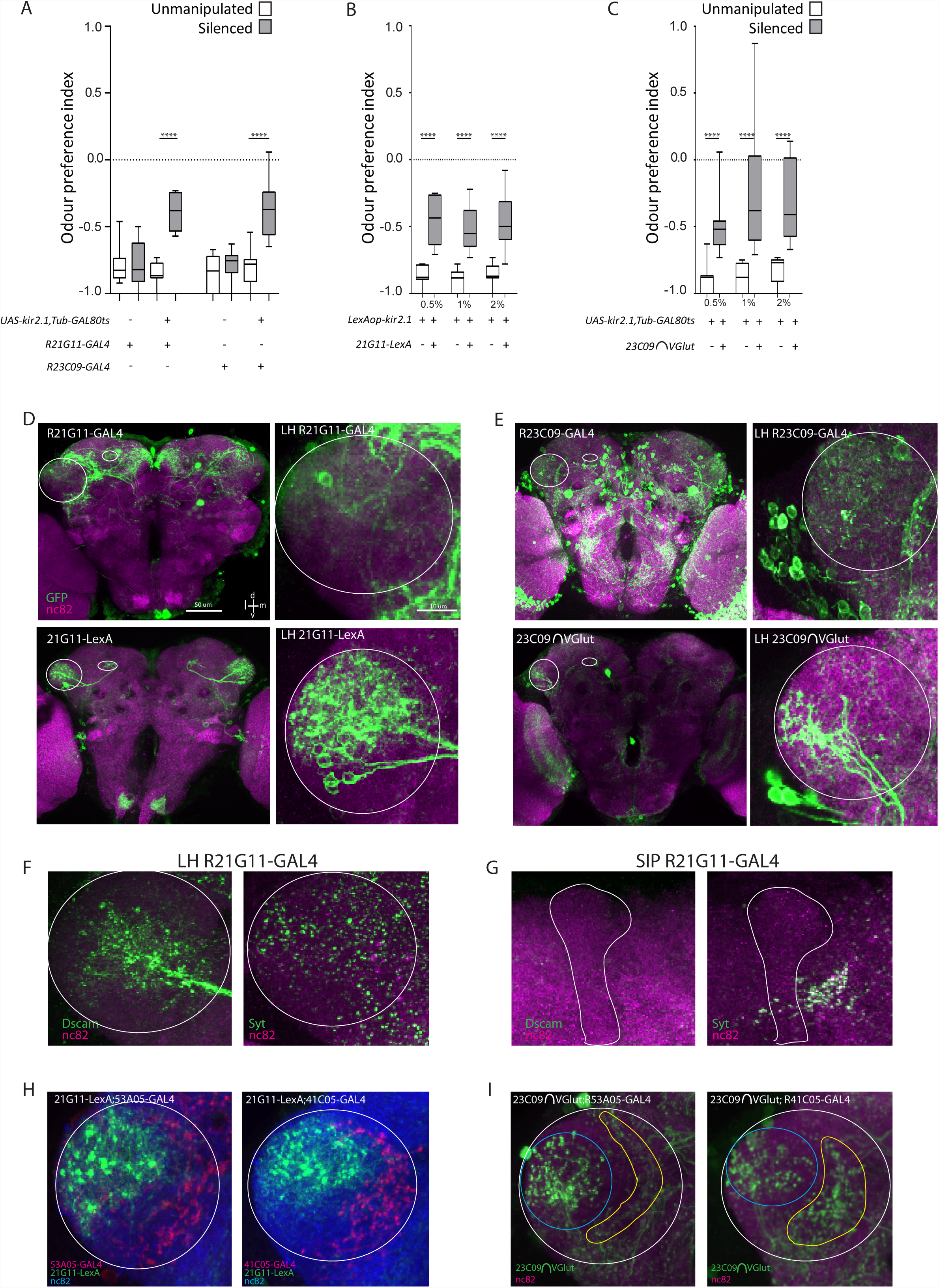
Activity in 21G11 and 23C09 neurons is required for behavioral response to CO_2_. (A) T-maze response to 1% CO_2_ of R21G11-GAL4 and R23C09-GAL4 flies driving UAS-Kir, TubGAL80TS expression. (A and C) White boxes, no heat induction of Kir2.1 expression (“unmanipulated”, see materials and methods). Grey boxes, heat induction of Kir2.1 expression before test (“silenced”). The box represents the first and the third quartiles and the whiskers the 10th and 90th percentiles. The line across the box is the median. (B and C) T-maze response to three CO_2_ concentrations - 0.5%, 1 % and 2% - of the flies with 21G11-LexA driving LexAopKir2.1 expression (B) and 23C09∩VGlut driving UAS-Kir, TubGAL80TS expression (C). For 21G11-LexA driving LexAopkir2.1 expression, white boxes represent parental controls and grey boxes represent constitutive Kir expression. For 23C09∩VGlut driving UAS-Kir, TubGAL80TS expression, white boxes represent no heat induction of Kir expression and grey boxes represent heat induction of Kir expression before test. Post-hoc two-way ANOVA showed no significance when comparing among expressions for both control and test samples. (D) Brain and lateral horn (LH) UAS-mCD8-GFP expression of R21G11-GAL4 and 21G11-LexA (green). (E) Brain and LH UAS-mCD8-GFP expression of R23C09-GAL4 and 23C09∩VGlut (green). (D-E) Circle highlights the LH and oval highlights the superior intermediate protocerebrum (SIP). (F and G) Dscam 17.1-GFP and syt-HA expression in the LH and SIP of R21G11-GAL4 (green). In (F) circle highlights the LH. In (G) the vertical lobe of the mushroom body is drawn to facilitate visualization of the adjacent SIP. (H) LH showing expression of 21G11-LexA (green) and the V-projection neuron lines R53A05-GAL4 (red) and R41C05- GAL4 (red). Circle highlights the LH. (I) LH expression of 23C09 Vglut (blue circles) and the V-projection neuron lines R53A05-GAL4 (yellow circle) and R41 C05-GAL4 (yellow circle). White circle highlights the LH. For all images the brain neuropil was stained with nc82 (magenta and blue). d=dorsal; l=lateral; m=medial; v=ventral. N=9-10; ±SEM ****p<0.0001. All p values are calculated with one-way ANOVA.

Given that CO_2_ avoidance is reduced but not abolished for either line we tested flies with both sets of neurons silenced (Figure S2B). We observe no change in the phenotype indicating that the two sets of neurons do not complement each other, i.e., the activity of these populations may not be independent to drive avoidance responses (Post-hoc two-way ANOVA comparing individual and combined expressions not significant both for control and test samples). It has been previously shown that different projection neurons of the V-glomerulus are required for the behavioral response to different CO_2_ concentrations (9). Therefore we tested whether the requirement of *21G11* and *23C09* neurons for avoidance to CO_2_ was concentration dependent. For this experiment, we used lines *21G11-LexA* and *23C09 ∩ VGlut* that have a restricted expression when compared to *R21G11* and *R23C09* lines (Figure 1D and E, see below). We observe that silencing the activity of these neurons reduces the avoidance behavior of the flies in a comparable manner across odor concentrations (Post-hoc two-way ANOVA comparing across odor concentrations not significant both for control and test samples). These results suggest that *21G11* and *23C09* neurons contribute to CO_2_ avoidance independently of concentration (within the range that does not engage the acid sensing response).

Anatomical inspection reveals that line 21G11 labels one cluster of neurons that innervate the dorsoposterior area of the LH and project to the superior intermediate protocerebrum (SIP) (Figure 1D). Some morphological and physiological types of LH output neurons have been described previously (17,28). The *21G11* cluster with its posterior cell bodies and projections to the SIP does not appear to correspond to the described morphological categories.

We generated a *LexA* version of the line to allow independent manipulation of the 21G11 and 23CO9 clusters of neurons. The *LexA* version of 21G11 is very sparse. Additionally, the number of neurons labeled in the LH cluster is smaller. We counted 10 cell bodies in the *LexA* version and 16 to 18 cell bodies in the *GAL4* version (n=5). When we overlaid the expression of both lines we found that 7 to 9 cells were specific to *21G11-GAL4*, three cells specific to *21G11-LexA* and seven cells are common to both lines (Figure S3, n=9). Nevertheless, activity in *21G11-LexA* neurons is necessary to elicit full CO_2_ avoidance (see Figure 1C). Line *23C09* labels more than one cluster of neurons at the LH (Figure 1E). To narrow down the expression of line *23C09*, we generated a *splitGAL4* version and then we intersected the expression with that of different neurotransmitter-*splitGAL4* lines (29,30). We found that a glutamatergic cluster located posteriorly is involved in the response (Figure 1C and E). This cluster, which we will call *23C09 ∩ VGlut*, has 8 to 10 cell bodies (n=3) with processes only within the LH. For expression of these lines in 10 µm sections across the brain see Figure S4. In order to determine the polarity of *21G11* neurons we used the neural compartment markers Dscam17.1-GFP(31), for dendrites and Synaptotagmin-HA (32) for presynaptic areas. Dscam17.1-GFP signal localized exclusively to the LH, which indicates that these neurons receive inputs there, presumably olfactory. The synaptotagmin-HA signal, on the other hand, is localized both to the LH and the SIP. To exclude the possibility that the *GAL4* cluster holds a mix population of neurons with different polarities we marked the more restricted *21G11-LexA* neurons and observed the same distribution of Synaptotagmin-HA (Figure S5). These results suggest that *21G11* neurons output both in the SIP and the LH. Finally, we asked if *21G11* or *23C09* contact projections from the V-glomerulus.

Three distinct projection neurons from the V glomerulus (VPN) at the antennal lobe innervate the medial border of the LH (9,10). We tested two VPNs for which there are lines available with a strong visible projection. We do not see clear overlap at the LH between the innervation of the VPNs and the innervation of the LH neurons we identified (Figure 1H and I). This observation together with the fact that the reduction in avoidance is not complete indicates that additional LH neurons are involved in the response.

### *21G11* and *23C09 ∩ VGlut* neurons respond to CO_2_ in different concentration dependent manners

Having demonstrated that *21G11* and *23C09* neurons are required for the behavioral response to CO_2_, we next addressed the physiological response of these neurons. We measured the changes in internal free calcium levels in *21G11* and *23C09 ∩ VGlut* neurons upon stimulation with CO_2_. For this we expressed the genetically encoded calcium indicator *GCaMP6m* (33) in *21G11* and *23C09 ∩VGlut* neurons and recorded the calcium dynamics in a live fly preparation at the two-photon microscope. The neurons labeled by *R21G11* respond to all concentrations of CO_2_ with the peak Δ F/F increasing from 0.5 to 1% CO_2_ (Figure 2A-B, Wilcoxon signed-rank test w=66.00, p=0.0011). The peak response to 1% and 2% are not significantly different though the length of the response appears to be larger at 2% (Wilcoxon signed-rank test w=38.00, p=0.0727). We also tested responses to CO_2_ in the subset of *21G11* neurons labeled by *21G11-LexA* (Figure 2C-D). To our surprise this subset responds only to 0.5% CO_2_ (Wilcoxon signed-rank test w=15.00, p<0.0001). This observation suggests that within the *21G11* cluster different neurons are sensitive to different concentrations of CO_2_. However, in Figure 1C we observed that when these same neurons are silenced, the behavioral response to different CO_2_ concentrations does not change. Based on these findings we speculate that as soon as the activity of the neurons responsive to the lowest concentration is compromised the behavioral output to all concentrations is affected. Recordings of *23C09 ∩ VGlut* neurons show they respond all CO_2_ concentrations tested.

**Figure 2.**
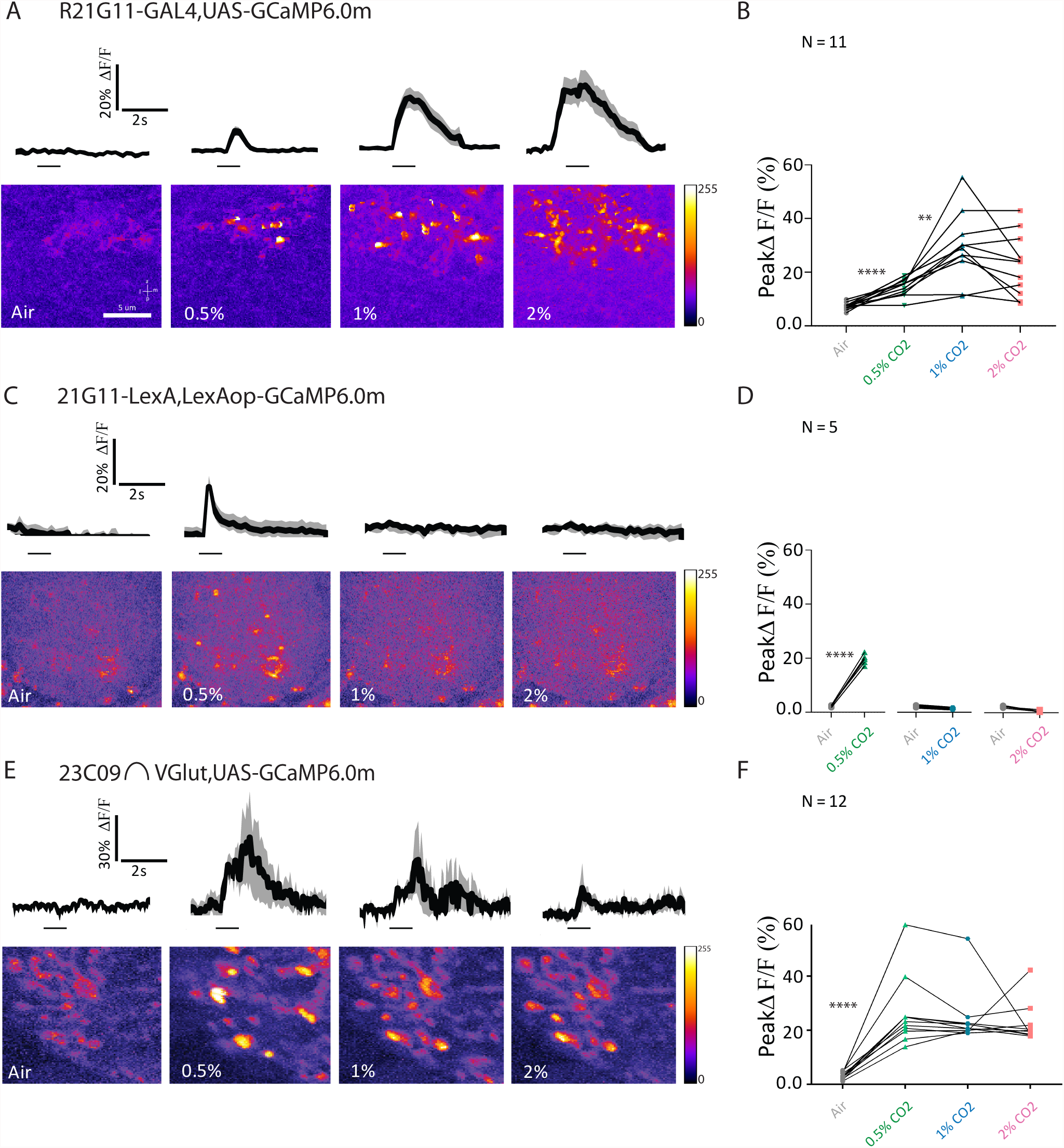
Physiological response to CO_2_ of 21G11 and 23C09 neurons. (A,C and E) LH activity of R21G11-GAL4, 21G11-LexA and 23C09∩VGlut, respectively to air, 0.5%, 1% and 2% of CO2. On top it is shown the average time course of GCaMP6.0m intensity change. The black bar indicates the time of the stimulus. The black trace represents the average while the grey shows the range of individual traces. On bottom the representative images showing the pseudo-colored response. (B,D and F) Peak GCaMP6.0m intensity change after stimulation with air, 0.5%, 1 % and 2% of CO2. a=anterior; l=lateral; m=medial; p=posterior. Error bars indicate ±SEM, **p<0.01,***p<0.001, ****p<0.0001. All p values are calculated with Wilcoxon signed-rank test.

Though the curve of the response appears larger for lower concentrations there is no significant difference between peak amplitudes of Δ F/F of different concentrations (Figure 2E-F). In summary, both clusters respond to CO_2_ stimulation at different concentrations, with each set of neurons exhibiting a different pattern of free calcium response to CO_2_ stimulation.

### Physiological response of *23C09 ∩ VGlut* depends on output of *21G11-LexA*

The two sets of neurons that we identified innervate a similar region of the LH and contribute to the same behavioral response. When we silenced both clusters simultaneously we saw no additive effect, indicating they are not independent from each other to drive avoidance (Figure S2B). We therefore asked whether the two clusters are connected. To this end we used GRASP (GFP reconstitution across synaptic partners) that reveals membrane contact between two sets of neurons (34). We observe a strong signal in the LH indicating that the membranes of the two clusters contact each other (Figure 3A). To assess functional connectivity we manipulated the activity of one cluster while imaging activity on the second cluster. We silenced *21G11-LexA* neurons with the expression of *Kir2.1* using the *LexA/LexAOp* expression system and imaged *23C09 ∩ VGlut* neurons expressing *GCaMP6.0m* with the *GAL4/UAS* system (Schematic in Figure 3B). To control for silencing we co-expressed *Kir2.1* and *GCaMP6.0m* in *21G11-LexA* neurons and confirmed that no calcium signal is observed with CO_2_ stimulation (Figure S6). Upon presentation of CO_2_ at the concentrations 0.5, 1 and 2%, we observed a very consistent response across trials and across concentrations in *23C09 ∩ VGlut* neurons when *21G11-LexA* neurons are silenced (Figure 3B and C, Mann-Whitney, not significant). When we compared the peak responses of *23C09 ∩ VGlut* while *21G11-LexA* is intact (Figure 2F) or silenced (Figure 3C), we found that there is pronounced reduction for 0.5% CO_2_ responses which corresponds to the profile of *21G11-LexA* responses (Figure 3D). The results indicate that the output of *21G11-LexA* neurons contributes to *23C09 ∩ VGlut* activity. To test this directly, we expressed the red-shifted channelrhodopsin *Chrimson* (35) in *21G11-LexA* neurons to allow activation of *21G11-LexA* neurons with 720nm light while recording *23C09n VGlut* calcium concentration with *GCaMP6.0m*. We observe that indeed activation of *21G11-LexA* neurons with light generates a strong calcium response in *23C09 ∩ VGlut* neurons (Figure 3E and F). No calcium response was observed in *23C09 ∩ VGlut* neurons when flies were not fed retinal, which is necessary for *Chrimson* function (Figure 3E and F).

**Figure 3.**
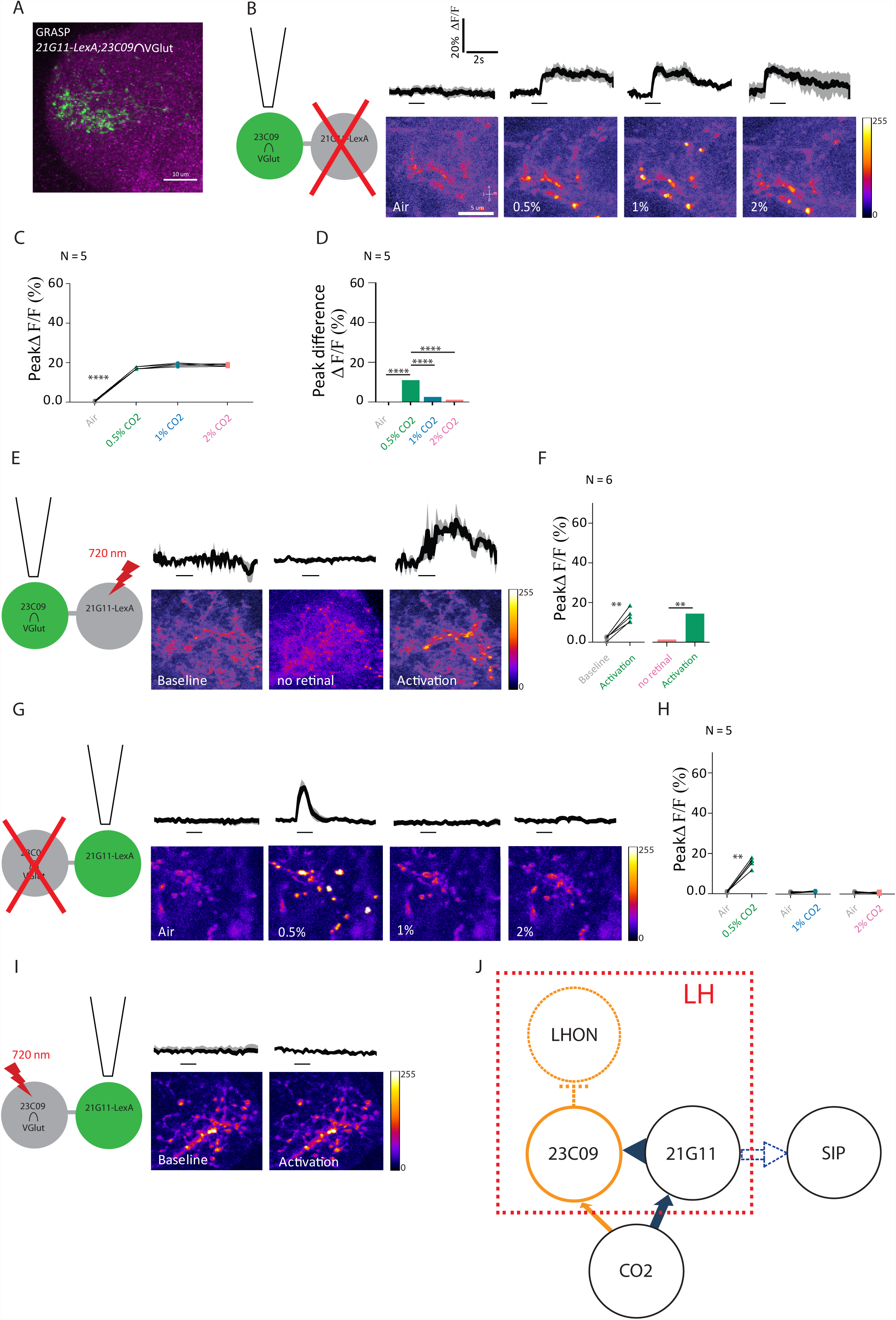
21G11 LexA neurons are pre-synaptic to 23C09 VGlut neurons. (A) Lateral horn showing GRASP, a measure of contact between the membranes of neurons, in 21G11-LexA and 23C09 VGlut. (B) Schematics of the experiment and calcium response at the LH, using GCaMP6m, of 23C09 VGlut neurons to air, 0.5%, 1 % and 2% of CO2 while 21G11-LexA neurons are silenced by expression of Kir2.1. (C and D) Peak GCaMP6.0m intensity change and peak difference in intensity after stimulation with air, 0.5%, 1 % and 2% of CO2. (E) Schematics of the experiment and calcium response at the LH of 23C09∩VGlut during baseline, activity without retinal and upon activation with 720 nm light of 21G11-LexA driving expression of Chrimson. (F) Peak GCaMP6.0m intensity change of 23C09 VGlut during baseline, activity without retinal and upon activation with 720 nm light of 21G11-LexA driving expression of Chrimson. (G) Schematics of the experiment and calcium response at the LH, using GCaMP6m, of 21G11-LexA neurons to air, 0.5%, 1 % and 2% of CO2 while 23C09∩VGlut neurons are silenced by expression of Kir2.1. (H) Peak GCaMP6.0m intensity change after stimulation with air, 0.5%, 1 % and 2% of CO2. (I) Schematics of the experiment and LH activity of 21G11-LexA upon activation of 23C09∩VGlut neurons, expressing Chrimson, with 720 nm light. (J) Proposed model of the LH neurons processing CO2 information. For (B), (E), (G) and (I), on top it is shown the average time course of GCaMP6.0m intensity change. The black bar indicates the time of the stimulus. On bottom the representative images showing the pseudo-colored response. a=anterior; l=lateral; m=medial; p=posterior. Error bars indicate ±SEM **p<0.01, ****p<0.0001. All p values are calculated with Wilcoxon signed-rank test.

We next did the converse experiments where we manipulate activity in *23C09 ∩ VGlut* neurons and image activity in *21G11-LexA* neurons using the same tools with the expression systems reversed. Silencing *23C09 ∩ VGlut* neurons does not change *21G11-LexA* response to CO_2_ presentation (Figure 3F and G). These results indicate that *23C09 ∩ VGlut* neurons do not output into *21G11-LexA* neurons. We then performed optogenetic activation of *23C09 ∩ VGlut* neurons and recorded the calcium response of *21G11-LexA* neurons. We did not observe a calcium response in *21G11-LexA* neurons upon light stimulation of *23C09 ∩ VGlut* neurons (Figure 3H). To control for activation of *23C09 ∩ VGlut* neurons we expressed both *Chrimson* and *GCaMP6m* in these neurons and could see a response with light stimulation (Figure S7). The activation results further support the notion that *23C09 ∩ VGlut* does not output into *21G11-LexA* neurons.

Taken together the results indicate that *21G11-LexA* neurons are presynaptic to *23C09 ∩ VGlut* neurons. The presence of a presynaptic marker at the LH processes of *21G11-LexA* neurons is consistent with these observations (Figure 1F). Based on our findings we propose a model where CO_2_ response is processed at the SIP via *21G11* output (Figure 3I). *21G11* also outputs at the LH to activate the *23C09 ∩ VGlut* local neurons that are glutamatergic. Given that it has been proposed that glutamatergic neurons in the fly brain are inhibitory (36), we speculate that *23C09* neurons inhibit output neurons that mediate attraction.

### *21G11* and *23C09 ∩ VGlut* neurons are selectively involved in processing CO_2_ avoidance

We then asked if the circuit we identified at the LH is involved in the response to other odors. It could be central to all odor responses or perhaps be involved only in the aversive responses or even selectively involved in the CO_2_ response. To answer this question we first measured the calcium responses to different odors (Figure 4 A-D). To test attractive odor responses we used farnesol, an attractant present in the rind of ripe citrus and processed through a single glomerulus (37), and apple cider vinegar, a complex attractive stimulus (38). While apple cider vinegar elicits a small response in both sets of neurons, farnesol does not elicit a response in either set of neurons (Figure 4A-D). To test aversive responses we used benzaldehyde that smells of bitter almond and acetic acid that elicits the acid sensing response in the antennal lobe (22). *21G11* neurons respond to both acetic acid and benzaldehyde (Figure 4A and B). *23C09* neurons respond only to benzaldehyde in an atypical fashion (Figure 4C-D). The rise in calcium concentration happens a few seconds after stimulus presentation. Though the peak ΔF/F in all these responses is low, the physiological response is broad and includes responses to aversive and attractive odors. How do these physiological responses translate into a behavioral response? To address this question we tested the requirement of activity in *23C09 ∩ VGlut* or *21G11-GAL4* (broader expression than *21G11-LexA*) neurons for the behavioral response to other odors. Similarly to what we did in the screen, we tested the flies using a T-maze. Also following the screen conditions, we silenced the neurons with *Kir2.1* only in the adult stage. We allowed the flies to choose between air and either farnesol, apple cider vinegar, acetic acid or benzaldehyde at 1/1000 dilution. It was reported that there is a non-olfactory component to benzaldehyde avoidance at 1/100 dilution. We confirmed that we are not including a non-olfactory component in our experiment by testing the response of flies without olfactory organs to our working dilution of benzaldehyde (Figure S8). We observe that activity in *23C09 ∩ VGlut* or *21G11-GAL4* neurons is not required either for attraction to farnesol or apple cider vinegar indicating that activity in these LH neurons is not involved in general odor responses (Figure 4E). Silencing *23C09∩VGlut* or *21G11-GAL4* neurons also does not affect avoidance to benzaldehyde and acetic acid (Figure 4F). Interestingly, though the arborization at the LH of the acid sensing projection neurons is very similar to the arborization of the V-glomerulus projection neurons (39), it appears that different LH neurons process these responses. The results indicate that activity in *21G11* and *23C09 ∩ VGlut* neurons is selectively required for the response to CO_2_.

**Figure 4.**
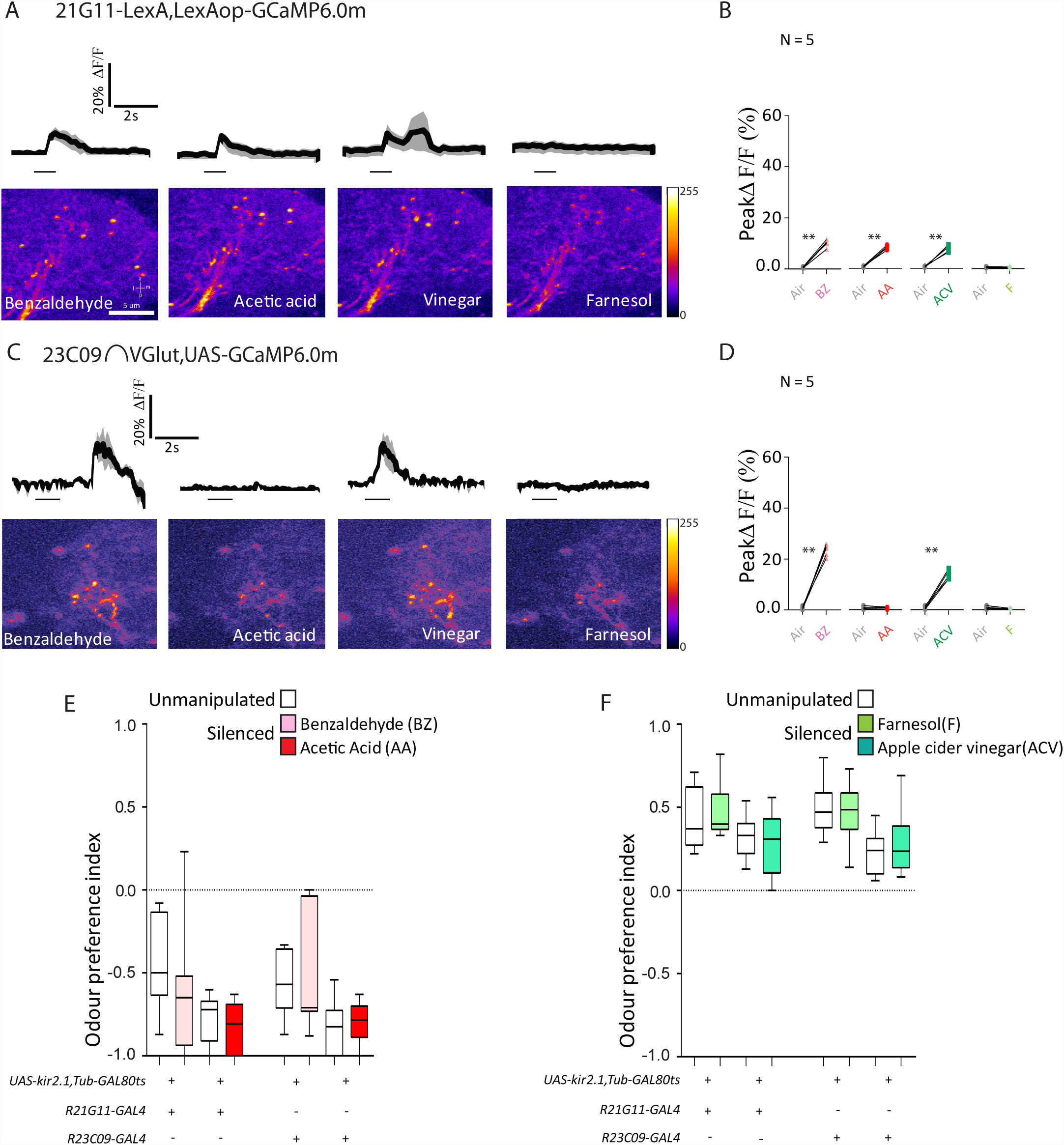
Physiological and behavioral response to attractive and repulsive compounds of R21G11 and R23C09 neurons. (A and C) Calcium response at the LH of 21G11-LexA and of 23C09 VGlut to the repulsive compounds – benzaldehyde (BZ) and acetic acid (AA)- and the attractive compounds – farnesol (F) and apple cider vinegar (ACV). On top it is shown the average time course of GCaMP6.0m intensity change. The black bar indicates the time of the stimulus. On bottom the representative images showing the pseudo-colored response. (B and D) Peak GCaMP6.0m intensity change after stimulation with benzaldehyde (BZ), acetic acid (AA), farnesol (F) and apple cider vinegar (ACV). (E) T-maze response to benzaldehyde (BZ) and acetic acid (AA) of R21G11-GAL4 and R23C09-GAL4. (F) T-maze response to farnesol (F) and apple cider vinegar (ACV) of 21G11-GAL4 and 23C09-GAL4. (E-F) White boxes, no heat induction of Kir2.1 expression (“unmanipulated”, see materials and methods). Colored boxes, heat induction of Kir2.1 expression before test (“silenced”). The box represents the first and the third quartiles and the whiskers the 10th and 90th percentiles. The line across the box is the median. N=5 for A, B, C and D; N=10 for E and F. For B and D Error bars indicate ±SEM **p<0.01, ****p<0.0001. p values are calculated with Wilcoxon signed-rank test. For E and F comparisons calculated with one-way ANOVA are non-significant.

## DISCUSSION

In this study we used a behavioral readout to directly address the role of the LH in olfactory responses. We showed that the activity of two clusters of LH neurons is required for *D. melanogaster* innate avoidance of CO_2_. Furthermore, because we have not found a correlation between the activity of these LH neurons and the behavioral responses to other general odors, we conclude that the activity of *21G11 and 23C09 ∩ VGlut* neurons is specifically required for responses to CO_2_. Thus, our results unravel a circuit within the LH region specific for CO_2_ avoidance responses in the lower concentration range. However, the avoidance to CO_2_ is not completely abolished by silencing these neurons. This could have three origins: not all cells are labeled in the identified cluster; unidentified clusters are involved; the LH has intricate connectivity and silencing of any LH neuron leads to disruption of the behavior. The latter is unlikely as we silenced and tested 32 lines labeling different LH neurons, some of them innervating very large sections of the LH and only the two lines described here consistently had an effect. Though we cannot eliminate any of the former possibilities, the lack of contact between the LH neurons we identified and the V-glomerulus projection neurons that carry CO_2_ stimulus information argues for the involvement of additional neurons.

Recent work indicates that MB can be involved in innate aversion and the LH is required for the execution of learned aversion (10,40–42). Hence, the roles of the two centers are not as segregated as previously thought and they must be connected to coordinate the innate and the learned responses. The *21G11* output neurons we identified connect to the superior intermediate protocerebrum. It is an area highly innervated by MB output neuron terminals (MBONs) suggesting a location for integration of LH and MB output (43,44). At the other extremity of *21G11* neurons, we showed that they output within the LH to the *23C09 ∩ VGlut* neurons. Since these are glutamatergic neurons and glutamatergic signaling in the brain can be inhibitory we speculate that activation of *21G11* and *23C09 ∩ VGlut* leads to inhibition of LH output neurons that mediate attraction.

A vast search has identified compounds that either increase or decrease the activity of the CO_2_ receptors (45). It was revealed that odors increasing receptor activity generate avoidance responses and odors decreasing receptor activity generate attraction responses indicating that the CO_2_ receptor pathway has a strong weight in establishing odor valence (46). Further work should elucidate how the LH neurons identified here contribute to these responses.

In summary, we demonstrated a role of the LH in an innate behavioral response. The neurons we identified appear to be involved in suppression of attraction at the LH. Moving forward, it will be interesting to explore whether a similar organization at the LH is used to generate responses to other odors and how the response is coordinated with the MB.

## ACKNOWLEDGEMENTS

We thank Eugenia Chiappe, Susana Lima, Marta Moita and members of the Vasconcelos Laboratory for discussion and comments on the manuscript. Barry Dickson provided fly stock and Gerald M. Rubin provided a promoter plasmid. We would like to thank the Fly and Molecular platforms of Champalimaud Centre for the Unknown (CCU) for generating the transgenic flies and scientific hardware and software platforms of CCU for support with the optogenetic activation experiments. This work was supported by Fundação Champalimaud, a grant from Fundação para Ciência e tecnologia (FCT) PTDC/BIA-BCM/104898/2008, FCT fellowship SFRH/BDP/77360/2011 to N.V.

## AUTHOR CONTRIBUTION

N.V. and M.L.V. conceived and designed the project. M.G. performed the behavioral screen. N.V. performed all other experiments with the technical assistance of S.D. M.L.V. provided guidance and wrote the paper with N.V.

## MATERIALS AND METHODS

### Contact for Reagent and Resource Sharing

Further information and requests for resources and reagents should be directed to and will be fulfilled by the lead contact Maria Luísa Vasconcelo.

### Experimental Model and Subject Details

Flies were maintained on standard cornmeal-agar medium at 18°C or 25°C and 70% relative humidity under a 12h light/dark cycle. Fly strains used were as follows: *UAS-Kir2.1* (26); *tub-GAL80^TS^* (27); *UAS-Chrimson* (35); *UASGCaMP6.0m* (33); *UAS-Dscam 17.1-GFP* (31); *UAS-syt-HA* (32); *UAS-CD4-GFP1-10* (34); *UAS-CD8-GFP* (47); *UAS-VGlutDBD* (30); *UAS-myr-tdTomato*(48);*LexAop-Kir2.1 (provived by Barry Dickson)* (49); *LexAop-Chrimson* (50); *LexAop GCaMP6.0m* (33); *LexAop-syt-HA* (51); *LexAop-CD4-GFP11* (34); *LexAop-mCD2-GFP*(52). We used the following *GAL4* lines: *R21G11*; *R23C09*; *R84A06*; *R65D12*; *R19B07*; *R13A11*; *R30A10*; *R37G11*; *R33E01*; *R93D02*; *R93D05*; *R41F11*; *R85C07*; *R64B02*; *R36E10*; *R30H02*; *R25B07*; *R36G09*; *R23F06*; *R54G12*; *R16C09*; *R13A07*; *R22B02*; *R29F04*; *R16C06*; *R82E01*; *R25A01*; *R84G12*; *R25G10*; *R26C12*; *R20C09*; *R20B0* from the Janelia farm collection (24,25).

## Genotypes per Figure

### Figure 1

#### Panel A

*w^1118^; UAS-Kir2.1, tub-Gal80^TS^; +*
*w^1118^; UAS-Kir2.1, tub-Gal80^TS^; 21G11-GAL4*
*w^1118^; UAS-Kir2.1, tub-Gal80^TS^; 23C09-GAL4*

#### Panel B

*w^1118^; LexAop-Kir2.1; +*
*w^1118^; LexAop-Kir2.1; 21G11-LexA*

#### Panel C

*w^1118^; UAS-Kir2.1, tub-Gal80^TS^; +*
*w^1118^;UAS-Kir2.1,tub-Gal80^TS^;23C09-AD/UAS-VGlut-DBD*

#### Panel D

*w^1118^; +; 21G11-GAL4/UAS-mCD8-GFP*
*w^1118^; LexAop-mCD2-GFP/+; 21G11-LexA/+*

#### Panel E

*w^1118^; +; 23C09-GAL4/UA S-mCD8-GFP/+*
*w^1118^; 23C09-AD/UAS-VGlut-DBD; UAS-mCD8-GFP/+*

#### Panel F and G

*w^1118^; UAS-Dscam-17.1-GFP; 21G11-GAL4 w^1118^; UAS-syt-HA; 21G11-GAL4*

#### Panel H

*w^1118^; 21G11-LexA/LexAop-CD2-GFP; UAS-myr-tdTomato/53A05-GAL4 w^1118^; 21G11-LexA/ LexAop-CD2-GFP; UAS-myr-tdTomato /41 C05-GAL4*

#### Panel I

*w^1118^; 23C09-AD/UAS-VGlut-DBD/+;UAS-mCD8-GFP/53A05-GAL4*
*w^1118^; 23C09-AD/UAS-VGlut-DBD/+; UAS-mCD8-GFP/41 C05-GAL4*

### Figure 2

#### Panel A and B

*w^1118^;21G11-GAL4; UAS-GCaMP6.0m*

#### Panel C and D

*w^1118^;21G11-LexA; 21G11-LexA/LexAop-GCaMP6.0m*

#### Panel E and F

*w^1118^; 23C09-AD/UAS-VGlut-DBD; UASGCaMP6.0m*

### Figure 3

#### Panel A

*w^1118^; LexAop-CD4-GFP11/21G11-LexA; UAS-mCD4-GFP1-10/23C09-GAL4*

#### Panel B, C and D

*w^1118^; 21G11-LexA, LexAop-Kir2.1/23C09-AD, UAS-VGlut-DBD; 21G11-LexA/UASGCaMP6.0m*

#### Panel E and F

*w^1118^; 21G11-LexA, LexAop-Chrimson/23C09-AD, UAS-VGlut-DBD; 21G11-LexA/UASGCaMP6.0m*

#### Panel G and H

*w^1118^; 21G11-LexA/23C09-AD, UAS-VGlut-DBD; 21G11-LexA, LexA opGCaMP6.0m/UAS-Kir2.1*

#### Panel I

*w^1118^; 21G11-LexA/23C09-AD, UAS-VGlut-DBD; 21G11-LexA, LexA opGCaMP6.0m/UAS-Chrimson*

### Figure 4

#### Panel A and B

*w^1118^;21G11-LexA; 21G11-LexA/LexAop-GCaMP6.0m*

#### Panel C and D

*w^1118^; 23C09-AD/UAS-VGlut-DBD; UASGCaMP6.0m*

#### Panel E and F

*w^1118^; UAS-Kir2.1, tub-Gal80^TS^; +*
*w^1118^; UAS-Kir2.1, tub-Gal80^TS^; 21G11-GAL4*
*w^1118^; UAS-Kir2.1, tub-Gal80^TS^; 23C09-GAL4*

### Figure Supp 1

#### Panel A

*w^1118^;UAS-Kir2.1, tub-Gal80^TS^; 84A06-GAL4*
*w^1118^;UAS-Kir2.1, tub-Gal80^TS^; 65D 12-GAL4*
*w^1118^;UAS-Kir2.1,tub-Gal80 ^TS^;21G11-GAL4*
*w^1118^;UAS-Kir2.1, tub-Gal80^TS^; 23C09-GAL4*
*w^1118^;UAS-Kir2.1,tub-Gal80^TS^;19B07-GAL4*
*w^1118^;UAS-Kir2.1,tub-Gal80^TS^;13A11-GAL4*
*w^1118^;UAS-Kir2.1, tub-Gal80^TS^; 30A 10-GAL4*
*w^1118^;UAS-Kir2.1, tub-Gal80^TS^; 37G11-GAL4*
*w^1118^;UAS-Kir2.1, tub-Gal80^TS^; 33E01-GAL4*
*w^1118^;UAS-Kir2.1, tub-Gal80^TS^; 93D02-GAL4*
*w^1118^;UAS-Kir2.1, tub-Gal80^TS^; 93D05-GAL4*
*w^1118^;UAS-Kir2.1,tub-Gal80 ^TS^;41F11-GAL4*
*w^1118^;UAS-Kir2.1, tub-Gal80^TS^; 85C07-GAL4*
*w^1118^;UAS-Kir2.1, tub-Gal80^TS^; 64B02-GAL4*
*w^1118^;UAS-Kir2.1, tub-Gal80^TS^; 36E10-GAL4*
*w^1118^;UAS-Kir2.1, tub-Gal80^TS^; 30H02-GAL4*
*w^1118^;UAS-Kir2.1, tub-Gal80^TS^; 25B07-GAL4*
*w^1118^;UAS-Kir2.1, tub-Gal80^TS^; 36G09-GAL4*
*w^1118^;UAS-Kir2.1,tub-Gal80^TS^;23F06 GAL4*
*w^1118^;UAS-Kir2.1,tub-Gal80 ^TS^;54G12-GAL4*
*w^1118^;UAS-Kir2.1,tub-Gal80 ^TS^;16C09-GAL4*
*w^1118^;UAS-Kir2.1,tub-Gal80^TS^;13A07-GAL4*
*w^1118^;UAS-Kir2.1, tub-Gal80^TS^; 22B02-GAL4*
*w^1118^;UAS-Kir2.1,tub-Gal80 ^TS^;29F04-GAL4*
*w^1118^;UAS-Kir2.1,tub-Gal80 ^TS^;16C06-GAL4*
*w^1118^;UAS-Kir2.1, tub-Gal80^TS^; 82E01-GAL4*
*w^1118^;UAS-Kir2.1, tub-Gal80^TS^; 25A01-GAL4*
*w^1118^;UAS-Kir2.1,tub-Gal80 ^TS^;84G12-GAL4*
*w^1118^;UAS-Kir2.1,tub-Gal80 ^TS^;25G10-GAL4*
*w^1118^;UAS-Kir2.1, tub-Gal80^TS^; 26C 12-GAL4*
*w^1118^;UAS-Kir2.1, tub-Gal80^TS^; 20C09-GAL4*
*w^1118^; UAS-Kir2.1, tub-Gal80^TS^; 20B07-GAL4*

#### Panel B

*w^1118^;UAS-Kir2.1, tub-Gal80^TS^; 65D 12-GAL4*
*w1118; UAS-Kir2.1, tub-Gal80^TS^; 21G11-GAL4*
*w^1118^;UAS-Kir2.1, tub-Gal80^TS^; 23C09-GAL4*
*w^1118^;UAS-Kir2.1, tub-Gal80^TS^;13A 11-GAL4*
*w^1118^; UAS-Kir2.1, tub-Gal80^TS^; 33E01-GAL4*
*w^1118^;UAS-Kir2.1, tub-Gal80^TS^; 85C07-GAL4*
*w^1118^; UAS-Kir2.1, tub-Gal80^TS^; 36G09-GAL4*
*w^1118^; UAS-Kir2.1, tub-Gal80^TS^;16C09-GAL4*

### Figure Supp 2

#### Panel A and B

*w1118; UAS-Kir2.1, tub-Gal80^TS^; +*
*w^1118^; UAS-Kir2.1, tub-Gal80^TS^; 21G11-GAL4*
*w^1118^; UAS-Kir2.1, tub-Gal80^TS^; 23C09-GAL4*

### Figure Supp 3

*w^1118^; LexAop-mCD2-GFP/+; 21G11-LexA/+*
*w^1118^; 23C09-AD/UAS-VGlut-DBD; UAS-mCD8-GFP/+*

### Figure Supp 4

*w^1118^;UAS-myR-TdTomato/LexAop-mCD2-GFP; 21G11-LexA/21G11-GAL4*

### Figure Supp 5

*w^1118^; UA S-syt-HA;21G11-LexA*

### Figure Supp 6

#### Panel A and B

*w^1118^;21G11-LexA, LexAop-Kir2.1/21G11-LexA/UAS-GCaMP6.0m*

#### Panel C and D

*w^1118^;23C09-AD/UAS-VGlut-DBD;UASGCaMP6.0m/UAS-Kir2.1*

### Figure Supp 7

#### Panel A and B

*w^1118^;21G11-LexA, LexAop-Chrimson/21G11-LexA/UAS-GCaMP6.0m*

#### Panel C and D

*w^1118^;23C09-AD/UAS-VGlut-DBD;UASGCaMP6.0m/UAS-Chrimson*

### Figure Supp 8

#### Panel A

*w^1118^; UAS-Kir2.1, tub-Gal80^TS^; +*

### Method Details

#### Generating transgenic flies

For the establishment of the *21G11LexA* line we used the Gateway System. The *LexA* vector used was purchased from Addgene (Plasmid #26230). We carried out a genomic PCR with the primers (5’ to 3’) F-GGGGACAAGTTTGTACAAAAAAGCAGGCTTCGCGCAGCACGTGAAGAACAAGGC and RGGGGACCACTTTGTACAAGAAAGCTGGGTCATGGCAACGTACTTCCAGTCCTCT. The fragment was inserted in a pDONR221 vector and then recombined into the pBPnlsLexA::p65Uw vector (Addgene, Plasmid #26230). To construct the *23C09AD* line we use the fragment in a pDON that was kindly provided by the Rubin Lab. We recombined it as described above.

#### Immunostaining

Adult fly brains were dissected, fixed and stained using standard protocols. Briefly, tissue was dissected in phosphate-buffered saline (PBS), fixed in 4% PFA in PBL (PBS and 0.12M Lysine) for 30 min at room temperature, washed 3x for 5 min in PBT (PBS and 0.5% Triton X-100) and blocked for 15 minutes in 10% normal goat serum in PBT (Sigma, cat# G9023). Samples were then incubated with primary antibodies for 72h at 4°C. After they were washed 3x for 5 min in PBT and incubated with secondary antibodies for 72h at 4°C. Finally the samples were washed 3x for 5 min in PBT and mounted in Vectashield (Vector laboratories, cat# H-1000). As primary antibodies we used: rabbit anti-GFP (1:2000, Molecular Probes, cat# 11122), chicken anti-GFP (1:2000, Molecular Probes, cat# A10262), rabitt anti-DsRed (1:2000, Molecular Probes, cat# 710530) and mouse anti-nc82 (1:10, Developmental Studies Hybridoma Bank). The secondary antibodies used were anti-rabbit or anti-chicken IgG conjugated to Alexa 488, anti-mouse or anti-rabitt IgG conjugated to Alexa 594 and anti-mouse IgG conjugated to Alexa 405. All microscopy of immunostainings was performed with a Zeiss LSM 710 confocal microscope. Images were processed with ImageJ.

#### Behavioral experiments

*Neuronal silencing* – Flies were kept at 18°C for 8 to 16 days. When using the *UASKir2.1,TubGAL80^ts^*, tester flies were placed at 30°C 24h before the experiment while control flies were always kept at 18°C. On the day of the experiment both tester and control flies were transferred to 25°C where we quantified their response on a T-maze (53). When using the *LexAopKir2.1* both tester and control flies were always kept at 25°C. We quantified flies response to air, three concentrations of CO_2_ (0.5%, 1% and 2%), two known attractive (apple cider vinegar (ACV) and farnesol (F)) and two known repulsive compounds (benzaldehyde (BZ) and acetic acid (AA)). To obtain the CO_2_ concentrations we mixed bottled synthetic air with bottled CO_2_ (Linde). The flow rate used was of 0.12 l per min. All other compounds were diluted 1:1000 in paraffin oil (Sigma). To test for the non-olfactory component to BZ avoidance the olfactory organs were removed manually (antennae and maxillary palps) 24h before the experiment. For all experiments flies were tested in groups of 20 individuals. Flies were placed on the T-maze elevator and dropped to the choice area where they were given 45 seconds to choose an arm. To control if the T-maze was balanced, we tested flies to air on both sides. For control and tester flies, one arm of the T-maze released air while the other arm released the testing compound. After the experiment, flies were counted and the odor preference index was calculated by subtracting the number of flies on the air side from the number of flies on the other side and dividing it with the total number of flies.

#### Calcium imaging experiments

##### Preparation

For all calcium imaging experiments flies expressed the calcium indicator *GCaMP6.0m*. The preparation was based on walking behavior preparation (54) but without the ball. To image the lateral horn (LH) in an *in* vivo preparation we glued the fly head to a microscope base (Scientifica) with bee’s wax (Sigma). We then opened a window that corresponded to half of the fly brain and removed all fat and trachea. We made sure that both antennae were untouched and healthy.

##### Microscopy

We used an Ultima two-photon laser-scanning microscope from Prairie Technologies (now Bruker) and a Coherent Chameleon XR lasers. All images were acquired every 0.2 ms with an Olympus BX61 microscope equipped with a 40×0.8 NA objective. To measure the fluorescent intensity at the LH we used ImageJ. The region of interest was delineated by hand and the result time trace was used for further analysis. To calculate the normalized change in the relative fluorescence intensity we used the Δ F/F=100(F_n_-F_0_)/F_0_ where F_n_ is the *nth* frame after stimulation and F_0_ is the average basal fluorescence before the stimulation. Images with visible rhythmic movements of the animal were discarded.

##### Olfactory stimulation

For olfactory stimulation a custom made delivery system consisting of a four-way solenoid valve (Parker Hannifin) connected to a peristaltic pump (Ismatec) creating a continuous airstream (1800 ml/min) that was delivered to the antennae with chemically inert tubing (Ismatec). The valve stimulation was commanded through the PrairieView software. For the CO_2_ stimulation, dilutions were placed in Tedlar gas sampling bags (#24634, Sigma) that were then connected to the valve. For stimulation with other compounds dilutions were made in glass vials with rubber taps (Fisher). At the rubber tap we inserted two venofix needles (Braun): one to connect the vial to the valve, the other to connect the vial to the air in the room for air-flow in the vial. We setup the system so that when a stimulus is triggered the odor dilutions replace only 50% of the air-flow in order to minimize the turbulence. For this reason all dilutions were prepared to double of the desired concentrations. In all experiments stimuli were delivered for one second. To control for calcium changes with the air-flow, all stimulations were repeated twice. In addition we performed experiments both in and outside the LH, and imaged the LH in neurons with both *Kir2.1* and *GCaMP6.0m*. No interference from the air-flow in the calcium response was ever observed.

##### Neuronal activation

For the neuronal activation we used 8 LEDs of 720 nm wavelength in a custom made ring placed beneath the stage that surrounded the fly. The delivery of light to the fly was commanded through the PrairieView software. The stimulus was delivered for one second at 5 Hz and 40 ms pulses.

##### Neuronal silencing

For the neuronal silencing experiments all flies expressed *Kir2.1*.

### Quantification and Statistical Analysis

All behavioral data was statistically analyzed by one-way analysis of variance and a Sidak’s multiple comparisons test. For all imaging data a Wilcoxon signed-rank test comparison was performed. For all analysis and statistical tests we used the GraphPad Prism Software version 6.0 (GraphPad Software).

